# figureComposer: A web-based interactive multi-panel bio-infographic designing tool

**DOI:** 10.1101/2020.03.04.976589

**Authors:** Kejie Li, Jessica Hurt, Christopher D. Whelan, Ravi Challa, Dongdong Lin, Baohong Zhang

## Abstract

**Background:** Many fit-for-purpose bioinformatics tools generate plots to investigate data quality and illustrate findings. However, assembling individual plots in different formats from various sources into one high-resolution figure requires mastery of commercial tools or even programming skills. In addition, it is a time-consuming and sometimes frustrating process even for scientists with modest computational skills.

**Results:** We developed figureComposer, a web-based bioinformatics tool that interactively arranges high-resolution images in various formats, mainly SVG to produce one multi-panel publication-quality composite figure in both PDF and interactive HTML formats in a user-friendly matter, requiring no programming skills.

**Conclusions:** figureComposer is open-source and publicly available web tool that can be accessed online at https://baohongz.github.io/figureComposer while the source code is provided at https://github.com/baohongz/figureComposer.

## Background

Scalable Vector Graphics (SVG) is an Extensible Markup Language (XML)-based vector image format, scalable to any resolutions without blurry pixelization as in other popular image formats, such as png, gif and jpg. This format has become one of the most broadly used image outputs adopted by many data analysis tools in computational biology field, notably R (1) and ggplot2 (2). In addition, SVG is usually set as the default image output by many JavaScript-based plotting libraries like D3 (3) and bioinformatics tools such as Coral (4), and rendered naturally by modern web browsers, such as Chrome, Firefox, Safari and MS Edge.

Composing multi-panel high resolution plots, such as those presented in Figure 1, usually poses a challenge for scientists with no or modest programming skills after gathering individual plots from various sources. User-friendly commercial tools such as Microsoft Power Point could be a viable option to arrange such plots, but these tools either cannot deal with complex pathway diagram in SVG format, or render this format in low resolution, sometimes even in malformed appearance as shown in Fig. s2.

**Figure 1.**
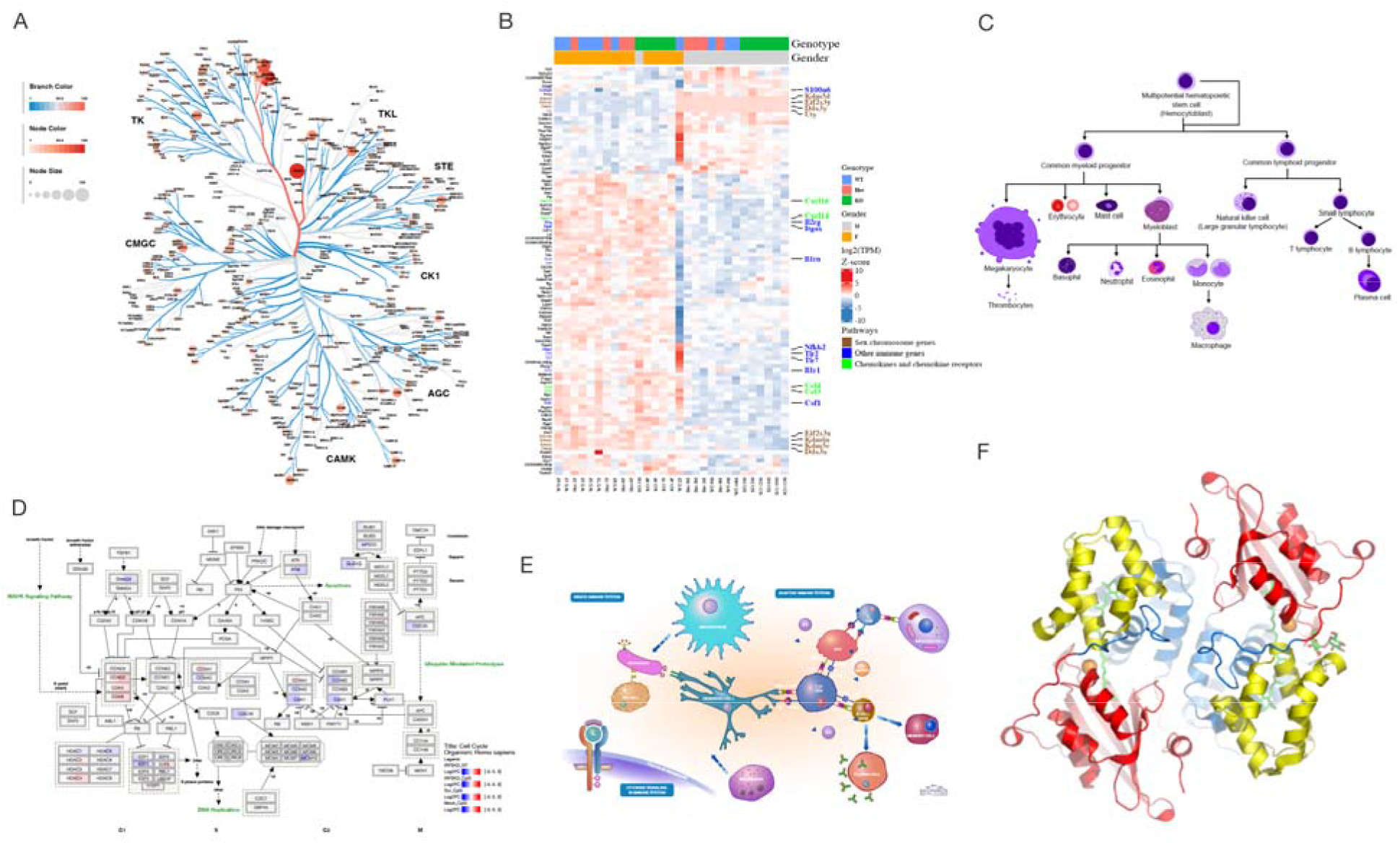
High-resolution plots generated by various tools are arranged by the online tool, figureComposer. Unless specified, the source plot is in SVG format. A) Human kinome tree generated by Coral web app (https://unc.live/2SqKj7s); B) Gene expression heatmap by R package ComplexHeatmap (8); C) Blood cell lineage from Wikimedia commons (https://bit.ly/2UPGTwx); D) Human cell cycle pathway diagram from WikiPathways (https://bit.ly/2SqRo8b); E) Human immune system illustration from Reactome (https://bit.ly/2Sp1ENX); F) Protein 3D structure ribbon form in png format by Pymol (https://pymol.org).

Continuing on the previous attempt of making a web-based interactive plot layout tool (5), we accept more image formats in figureComposer (Fig. s1) beyond only SVG as acquiring such format might be unfeasible in certain circumstances such as scanning gel images, and improved usability tremendously by implementing advanced functions outlined in the following section.

## Implementation

### Flexible text input

Regular characters plus built-in Emoji and symbols from Chrome browser can be used in the title of a plot, which is fully formattable in various font families, styles, sizes, shades, and colors by using an integrated text editor. Moreover, resizable boxes can be placed freely on the canvas to input paragraphs of text (Fig. s3).

### Versatile image formats

Besides SVG format, figureComposer takes other popular image formats including png, gif and jpg as input. For uncovered formats like tiff, free tools such as Inkscape (https://inkscape.org) (6) or pdf2svg (https://bit.ly/2NVtj6E) can be used to convert these to one of acceptable formats, preferably SVG.

### Stylesheets conflict

Since the scope of stylesheet definition used to style elements in SVG file is always global, which causes cross contamination of formatting when multiple SVG files are put together on the design canvas (Fig. s4). To overcome the shortcoming, the tool automatically converts these definitions into inline styles embedded to each targeting element individually and stores it locally before removing from globally scoped stylesheets.

### Image transparency

White background in SVG file is optionally removable to make it transparent so that plots can be overlaid onto each other to create compelling art as illustrated in Fig. s5.

### Vertical stacking

Each image is attached with a vertically stacked control button. Desired vertical stacking order is attainable by moving these control buttons up or down (Fig. s6) that provides additional dimension for creative design requiring overlaid images.

### Interactive HTML output

The finished work can be saved as a self-contained HTML file with necessary JavaScript code embedded for easy sharing by email or hosting at GitHub like services as exemplified at https://bit.ly/2IKu7lv.

## Results

We developed figureComposer, an interactive web-based bioinformatics tool that arranges high-resolution images in various formats, mainly SVG to produce one multi-panel publication-quality composite figure in both PDF and interactive HTML formats in a user-friendly manner, requiring no programming skills.

We compared it with several popular tools to illustrate the advanced features of figureComposer. Among six tools, except patchwork (7) that is command line based, the rest offers interactive user-friendly interface. In addition, figureComposer and canvasDesigner are conveniently accessible web-based tools. Regarding image format, Adobe Acrobat and patchwork won’t take SVG as input natively while Powerpoint and Inkscape have issues when rendering complex pathway diagrams in SVG format as displayed in Fig. s2. Although canvasDesigner and figureComposer have many common features, figureComposer breaks limitation of canvasDesigner by solving conflicting stylesheets problem, accepting images in various formats, overlaying images in any order vertically, and providing flexible text input. In summary, We list comparison scorecard of features among these tools including both open source solutions and popular commercial tools available to the authors in Table 1.

**Table 1.** Comparison scorecard of figure design tools

|  | figureComposer<br>v1.0 | canvasDesigner<br>v1.0 | MS Powerpoint<br>v16.3 | Adobe Acrobat<br>Pro DC v2020.006 | patchwork<br>v1.0 | Inkscape<br>v0.92 |
| --- | --- | --- | --- | --- | --- | --- |
| **Open source** | Yes | Yes | No | No | Yes | Yes |
| **Availability** | Free | Free | License fee | License fee | Free | Free |
| **Multi-image formats** | Yes | No | Yes | Yes | No | Yes |
| **Rendering speed** | Fast | Fast | Fast | Fast | Fast | Slow |
| **Text input** | Yes | No | Yes | Yes | Yes | Yes |
| **Interactive HTML output** | Yes | Yes | No | No | No | No |
| **SVG input** | Yes | Yes | Yes | No | No | Yes |

## Conclusions

figureComposer is an open-source and publicly available web-based tool that can be accessed online at https://baohongz.github.io/figureComposer while the source code is provided at https://github.com/baohongz/figureComposer. It has the most feasible features to improve productivity in the case of creating high-resolution multi-panel figures for scientific publications. Furthermore, the innovative HTML output brings a new way of illustrating high resolution figures interactively with unlimited zoom-in capability, which could be a nice feature for journals to incorporate in online publishing.

## Availability and requirement

Project name: figureComposer

Project home page: https://github.com/baohongz/figureComposer

Operating system(s): Platform independent

Programming languages: JavaScript

Other requirements: Internet connection and modern JavaScript-enabled web browsers, such as Chrome, Firefox, Safari and MS Edge.

License: GNU GPL version 3

## List of abbreviations

SVG: Scalable Vector Graphics;
XML: Extensible Markup Language;

## Declarations

### Ethics approval and consent to participate

Not applicable.

### Consent for publication

Not applicable.

### Availability of data and materials

All code, sample images and supplementary files can be found at project’s Github site https://github.com/baohongz/figureComposer https://baohongz.github.io/figureComposer/figureComposer_supplementary.pdf

### Authors’ contributions

KJ and BZ conceived and designed the tool that BZ implemented. All authors tested the tool and contributed to writing of the manuscript. All authors approved the final manuscript.

## Acknowledgements

Not applicable

